# Assessing predictive accuracy of species abundance models in dynamic systems

**DOI:** 10.1101/2025.01.23.634630

**Authors:** Christopher J. Brown, Christina Buelow, Rick D. Stuart-Smith, Neville Barrett, Graham Edgar, Elizabeth Oh

## Abstract

Cascading human pressures and environmental change are affecting the natural dynamics of animal populations. Forecasting population abundances from time-series data provides an important avenue for testing competing ecological theories, and for supporting conservation planning and sustainable use, yet changing system dynamics may lead to erroneous predictions. Predictions from a model fitted and tested on historical system dynamics may become irrelevant if system dynamics change. Here we describe methods to test forecast skill in a rapidly changing system where model parameters are likely to be non-stationary. We presented two ways to split time-series into training and test datasets so that training data were: 1) contemporary to the testing data (‘modern split’), and 2) not contemporary to the testing data (‘legacy split’). As a case-study, we use animal abundance data from a 30-year time-series from a global warming hotspot. We tested our approach on four temperate reef species with different temporal trends. The case-study and simulation tests confirmed larger forecast errors in legacy split when compared to the modern split. We found that the legacy split had errors that could be more than for times larger for a species that had a rapid collapse in abundance and non-stationary population dynamics. As expected for the species with rapid collapse, the legacy split estimated much higher forecast error than the modern split. Our approach is applicable to a large range of species and systems, including fisheries and threatened species population monitoring, where rapidly changing environments present threats to both the species and management efficacy. Accumulated lessons from across species and systems should shed light on critical generalities that precede broader ecosystem change.

## Introduction

Ecosystems are changing faster than has ever been observed previously by modern science. The pace of climate change is driving rapid ecological transformation (Poloczanska et al. 2013; Pecl et al. 2017). Many of these changes have been anticipated, however, climate change is accelerating more rapidly than expected and the scale of impacts is creating ecological responses that cannot be anticipated from historical analogues that are smaller in extent (Driscoll et al. 2024; Hughes et al. 2018). Climate impacts are also interacting with a multitude of other human-driven environmental changes (Côté, Darling, and Brown 2016). For instance, forest ecosystems may face tipping points of hydrological processes and extinctions as they become increasingly fragmented (e.g. Galán-Acedo et al. 2023; Xu et al. 2022). Human development of the oceans is accelerating, forcing change that has no historical analogues either in terms of the types of industries, like fish farming, or extent of existing industries, like fishing (Jouffray et al. 2020). There is also positive change for ecosystems; restoration efforts accelerate and create landscape scale benefits for ecosystems (e.g. Sievers et al. 2025).

A challenge for ecological science is developing models that can anticipate change on management relevant timescales of 1-10 years ahead (here referred to as ‘near-term’). Improving our ability to forecast ecosystems is an imperative because the rapid pace of human impacts is leading to unexpected ecological change on near-term timescales: we need our models to predict to environments that have no analogue in historical data. Anticipating this change could strengthen ecological theories (Lewis et al. 2023) and support adaptative management (e.g. Bradford et al. 2018; Liu et al. 2018; Henden et al. 2020). Yet, the rapid pace of environmental change also makes forecasting more difficult (Coron et al. 2012) because models fitted to historical data may therefore not represent future system dynamics – an issue often referred to in the time-series modelling literature as the technical issue of non-stationary model parameters (e.g. Cerqueira, Torgo, and Mozetič 2020). In spatial modelling literature, the match between the environmental parameter combinations that models are trained on and the conditions where we want to predict to has been conceptualized as model transferability (Yates et al. 2018). Improving models so they predict accurately to unprecedented environmental conditions and transfer well to different locations are important goals (Roberts et al. 2017; Yates et al. 2018), but there are ecological limits on how much accuracy can be gained with improved data and models (Dietze 2017; Beckage, Gross, and Kauffman 2011). Quantifying forecast error is important for management and planning, such as fishery management, which utilise well calibrated errors to infer the probability of fishery collapse (Raftery 2016). Likewise, limits in forecast accuracy can reveal the inadequacy of current ecological theories (Lewis et al. 2023).

Robust testing of forecasts should be done via validation or cross-validation, that is, testing models on data they were not trained with (Tashman 2000; Simonis, White, and Ernest 2021; Record, Boettiger, and Rollinson 2023). The common approaches for cross-validation of time-series models implicitly update training data so that it is contemporary to the testing data, therefore they may not reveal the degradation of forecast accuracy that is caused by no analogue futures. In the typical validation approach, the data are split in training and testing sets in multiple ways, to provide multiple estimates of forecast error (Tashman 2000). The decision rule for splitting is termed a ‘split-test rule’ and a common split-test rule is the rolling origin. For example, we would train the model on data from 1992-2006, the forecast origin then becomes 2006 and we test our model on 2007-2022. Then we roll the origin forwards one year, training on 1993-2007 and testing on 2008-2022. The established convention for addressing non-stationarity would then to explicitly account for it in the model’s structure when training (e.g. Ross et al. 2015). This convention treats non-stationarity as a technical issue that can be overcome with appropriate models and sufficient data. But for many ecosystems it is becoming increasingly difficult to find snalogues of future environments in historical data (Williams, Jackson, and Kutzbach 2007)—non-stationarity of system dynamics will be a persistent challenge for forecasting. As well as seeking to anticipate non-stationarity we need to quantify how predicting to futures with no analogue in the past will impact our estimation of errors. This issue of increasingly novel climates impacting forecast accuracy has been recognized in other disciplines, such as hydrology (Coron et al. 2012). It is likely that our ecological models will also be over-confident about the future environments with no analogue in past data (errors too low).

We propose a new split-test rule for quantifying forecast skill in dynamic ecological systems, which is to quantify how model accuracy changes when predicting into no analogue futures. In doing so, we acknowledge that no amount of data and modelling for historical ecosystems will ever generate a model that has errors that are well calibrated for future systems. Therefore, we should be quantifying how model accuracy deteriorates. Doing so may reveal new insights into the limits of accuracy of ecological forecasting, such as what types of species or ecosystems will be the hardest to predict. We approach the problem by developing a new split-test rule, which was designed to ensure that the training data are not contemporary to the testing data.

Our new splitting rule, which we term the legacy split, deliberately trains the model on data that is not contemporary to the testing sets and is inspired by similar efforts made for hydrological models (Coron et al. 2012). It allows for comparison to the conventional rolling origin splitting rules (which we here term ‘modern splits’) where the model is trained on data that is contemporary to testing data (Tashman 2000). This comparison means we can make inferences about how forecast accuracy may decline in systems where underlying dynamics change. We first confirmed the two splitting rules behaved as expected for simulated data. Then we tested the splitting rules on real time-series. We chose four species from a rapidly warming reef community that are exemplars of different types of change. We hypothesized that: 1. Forecast accuracy would decay for predictions further from the origin, i.e. at longer horizons (Petchey et al. 2015). 2. Forecast accuracy as evaluated with the legacy splitting rule for years that were further from the training period would be lower than forecast accuracy evaluated with the modern splitting rule. 3. The comparison of the legacy splitting rule to the modern rule would show the biggest difference for species that had rapid changes in abundance, whereas the two splitting rules would give similar results for species that have little or gradual long-term change.

## Materials and Methods

We followed the typical workflow for splitting time-series into training and test series to evaluate forecast accuracy: (1) Define the minimum training period; (2) split the timeseries into multiple training-test splits (Fig 2a); (3) fit model(s) to the training periods of each split; (4) calculate error statistics for each split for each forecasted year; (5) aggregate error statistics across splits for each forecast horizon (i.e. years forward from the forecast origin). Using multiple training-test splits reduces errors associated with unique windows of the timeseries and provides multiple tests at each forecast horizon. By summarizing over the forecast errors from all splits at each forecast horizon, we get more reliable estimates of model error.

**Figure 1.**
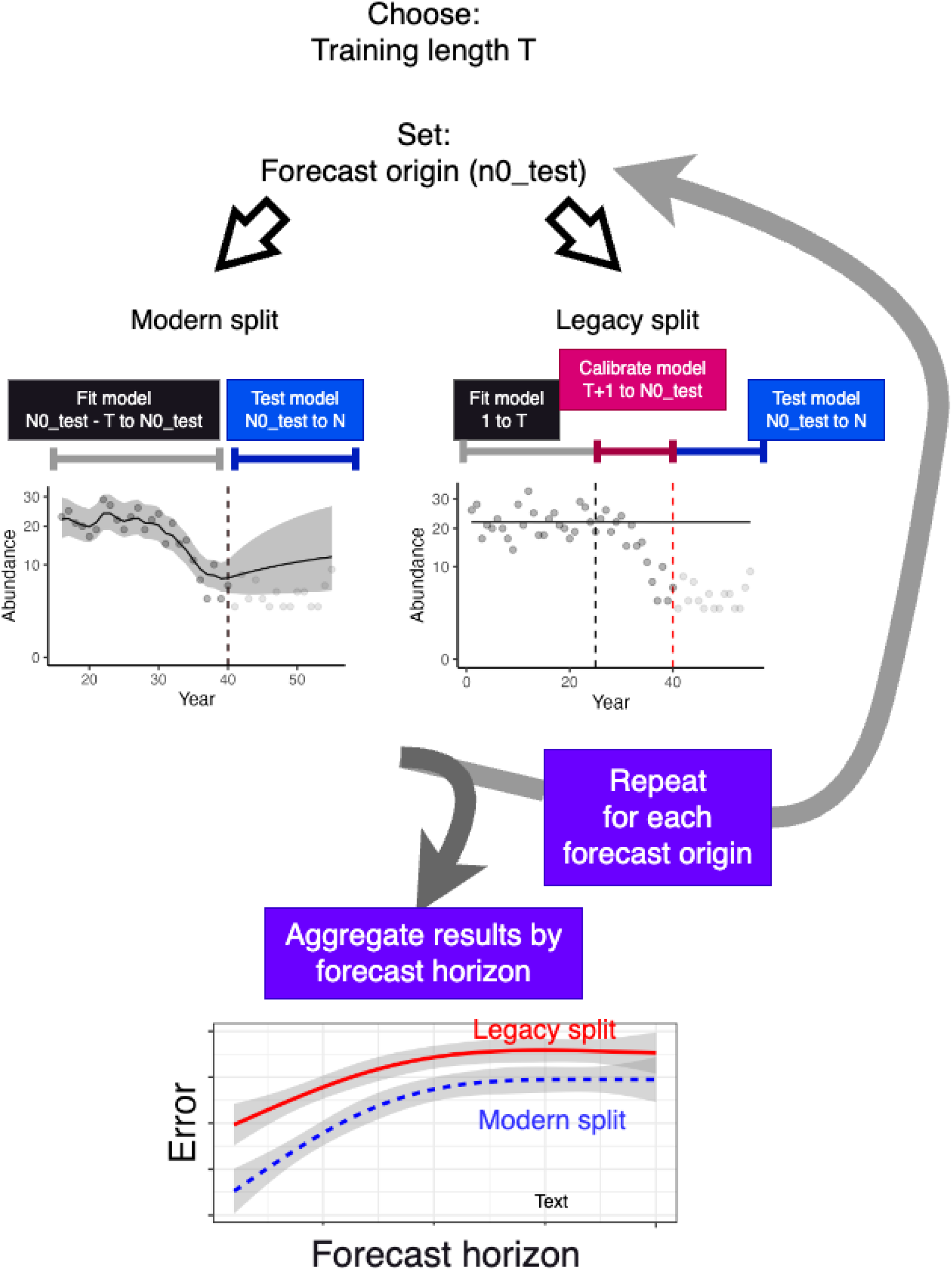
Workflow for modelling, include example plots of time-series fits and forecasts. Include conceptual diagram of hypotheses. Time-series length is chosen to give a model fit of sufficient quality for the study.

**Figure 2.**
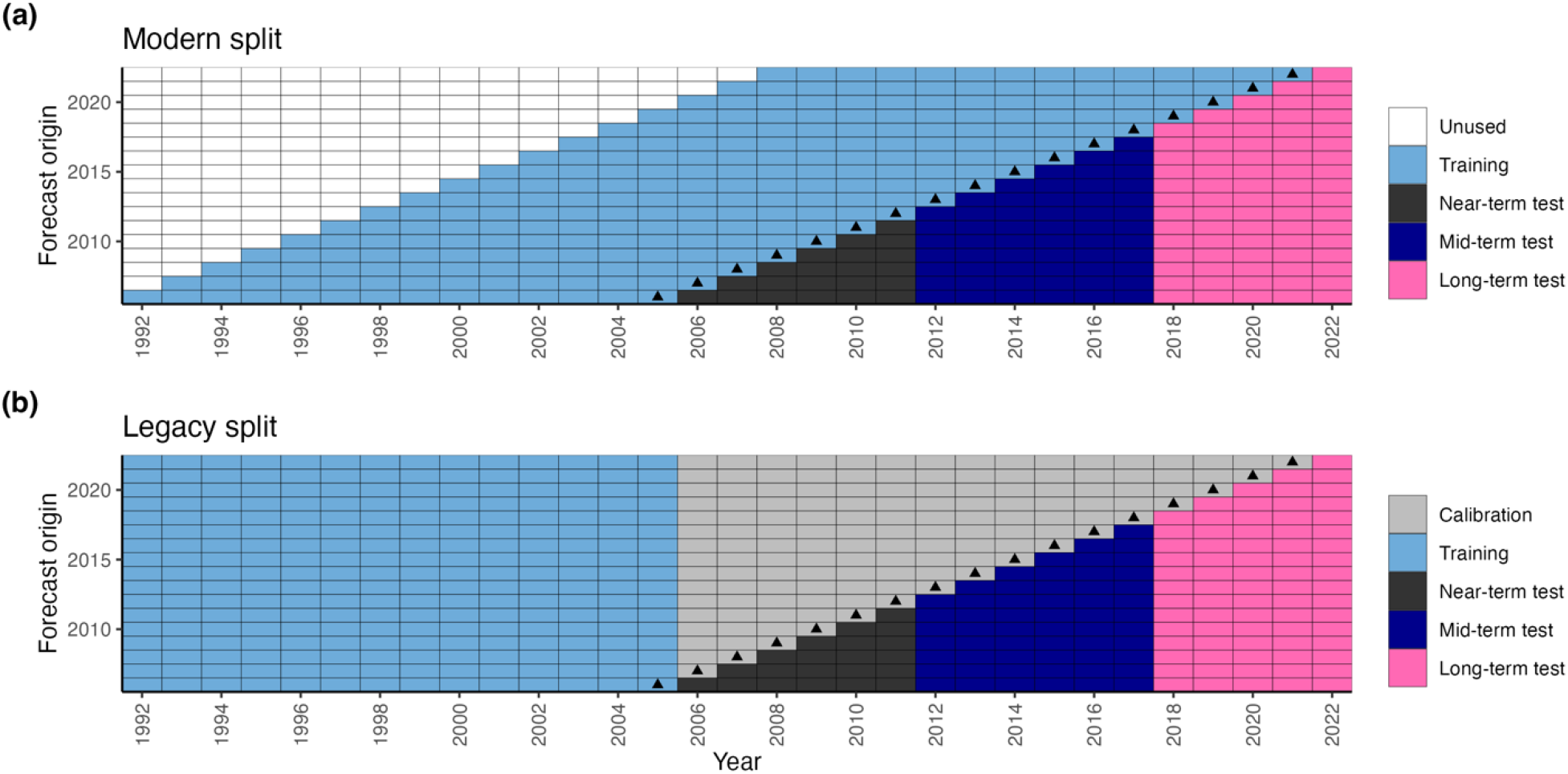
Modern splitting method with a rolling training period (a) and the legacy splitting method with a fixed training period (b) as applied to the Maria Island case-study dataset. Test data was divided into three epochs (near, mid and long-term) for analysis of model errors. The dot indicates the forecast origin.

### Split-test forecast rules

The core of our analysis is a modification of the most commonly applied splitting rules. We designed two rules to splitting a timeseries into multiple training and test sets (Figure 2). The first rule is common in the timeseries modelling literature (Tashman 2000; Simonis, White, and Ernest 2021). We developed the second rule to represent a case where parameters are not updated to new environments, thereby representing the situation we face when trying to predict future environments that have no analogue in contemporary data.

#### Modern split: Rolling training period and origin(Tashman 2000)

Iteratively train the model to a training period, then forecast for all future years, rolling forwards the initial and final years of training and the forecast origin for each test set (e.g. train 1992:2005, test 2006:2021, then train 1993:2006 test 2007:2021s etc…). This rule standardizes timeseries length in the training data, so we could compare whether model forecast skill changes through time, without confounding with the length of the training period. The downside to this method was that the model was increasingly trained on more contemporary data, so it may not provide a realistic estimation of future uncertainty where system dynamics may be considerably different. Pseudo-code for the algorithm:

1. for n0_train in 1:(N - T)
  1. Set nt_train = n0_train + T-1
  2. Train the model on data for n0_train:nt_train
  3. Set n0_test = nt_train
  4. Forecast and test for (n0_test+1):N Where: n0_train was the training origin n0_test is the forecast origin, N was the time-series length and T was the training time-series length.

#### Legacy split: Fixed training period, rolling origin

In this approach the model was trained once to an initial time-period. We then iteratively tested the model by rolling forwards the forecast origin (e.g. train on 1992:2005, then forecast for 2006:2021, 2007:2021 etc…). This rule meant that each new test period is increasingly further from the training period and therefore we expected it to provide error estimates that were representative of future forecasts where system dynamics have changed. To ensure that error estimates in this method were not conflated by changes in the initial condition for each split, we recalibrated the model up until the forecast origin. This recalibration initialized the mean predicted abundance to a new origin without updating the model parameters. Pseudo-code for the algorithm:

1. Train a model ‘m_train’ for data 1:nt_train
2. for n0_test in (nt_train+1):(N-1)
  1. Recalibrate the model over 1:n0_test with the parameters fixed at their values from m_train.
  2. Forecast and test for (n0_test+1):N

Note that a third method is also common in the literature. This is rolling origin evaluation (Tashman 2000) and is similar to (1) above except that training years are added to the initial training set, so that the length of the training series is increased with each iteration. We did not use that rule here because it does not standardize for length of training series.

The rolling origin approach, termed the ‘modern split method’ here, was originally proposed by Tashman (2000) who stated “Re-calibration, moreover, desensitizes error measures to events unique to the original fit period.” We developed rule (2) to ensure that error measures were sensitive to events unique to the original fit period. The one-off fitting meant this method is representative of a case where the future may have changed so much that the model parameters are now different.

### Time-series models

For both the simulation testing and the case-study we fitted Bayesian time-series models with the R-INLA program ((Rue et al. 2017) version 24.06.27, details in supplemental methods). For the case-study we fitted a series of models with Poisson observations and the linear predictor:

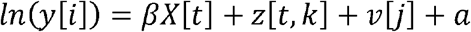

Where *y*[*i*] was the expected abundance for an individual survey *i, β* were coefficients for the covariates *X* [*t*], *z* [*t,k*] were the timeseries components for each region, *v* [*j*] were site specific random intercepts and a was a global intercept term. The simulation testing used the same model, but without covariates or site random intercepts.

We tested three common models for *z*[*t*], each representing different assumptions about the underlying population dynamics and different default forecasts (Table S1).

The first model was a random walk. Forecasts made by the random walk model have a mean that is the continuation of abundance at a constant level. The process-based interpretation of the random walk model is that each site has an independent population, where the standard deviation of the random walk equals the standard deviation in the intrinsic population growth rate (Brown and Roff 2019). This model assumes there was no density dependence.

The second model was an auto-regressive order 1 (AR1). The AR1 on a log-scale is equivalent to a Gompertz population growth model, and can be implemented for count data with environmental covariates in the R-INLA software (Ross et al. 2015). The AR1 converges on a mean at the Gompertz carrying capacity and the sum of its intercepts and environmental covariate effects is the intrinsic population growth rate (Ross et al. 2015). In the case-study, the four regions shared the priors and variance and autocorrelation parameters—meaning we assumed the different regions had the same Gompertz growth and carrying capacity parameters. We could thereby borrow strength across regions to estimate population parameters. Sharing parameter estimation was advantageous, because fished populations inside the protected area are likely to be closer to their carrying capacity, whereas the depleted abundances from outside the protected area may be insufficient to estimate a carrying capacity parameter.

The final model was a random walk order 2 (RW2). This model has no interpretation as a population dynamics model, but was included because it forecasts a continuation of trend.

### Validation statistic

Forecast skill was assessed with the median absolute scaled error (MASE). The advantage of the MASE is that it can be compared across different survey times and species (Ward et al. 2014; Tashman 2000) and is scale independent. The median over the mean is preferred when there are zeros in the data (Tashman 2000). The ASE was calculated:

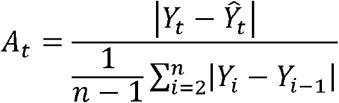

Where, *Y*_*t*_was the observed abundance at a time *t* that is part of the test 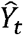 set, is the predicted abundance and *n* is the length of the training set. The data *Y*_*i*_, *i* ∈ 1: *n* is the training data. The absolute error (numerator) was scaled by the mean error in the training data (denominator). *A*_*t*_ was calculated for each survey site in each test year, and then the MASE was the median across all survey sites. The MASE converges to a value of 1 for long-timeseries that are a random walk.

Where the ASE (denominator) was zero, due to consistent zero abundances historically, we set it to a small number to represent lack of historical variance (=1e-4). This did result in some extreme MASE values, so to normalize results for plotting and analysis we set every MASE >90th quantile to the value of the 90th quantile.

### Simulation testing

We used a simulation test to verify that the legacy rule provided larger error estimates than the modern rule. We simulated time-series that were representative of common ecological time-series. The base case was one abundance survey per year for 55 years, with poisson observations and a log-link function. Population dynamics followed a Gompertz logistic growth model with log-normal process errors, such that the population size varies around the expected level according to the standard deviation of the random deviations. The environmental covariate was simulated as a lag-one autoregressive process, so that its timeseries had characteristics of low-period climate oscillations (e.g. Walters and Parma 1996).

To simulate a change in underlying system dynamics, we reduced the Gompertz carrying capacity parameter at year 32 by multiples of 0.25, 0.5 and 0.75. This reduction is indicative of regime shifts, such as sudden unexpected changes in fish stock productivity (Vert-pre et al. 2013) (full parameters in Table S2).

The initial training period for the time-series model was taken as years 1:25 (before the productivity shift) and then the training data was updated as per our two split-test rules. We summarized results by plotting mean forecast error for each horizon, plotting the errors by the two split-test rules before the collapse and +3 and +12 years after the collapse (Fig 2).

### Case-study

We used data from long-term monitoring of reef species at Maria Island, East Coast of Tasmania, Australia (Edgar and Barrett 2012; Edgar and Stuart-Smith 2014). This monitoring used a standardized dive survey method to survey 12 reefs over the period 1992 - 2021 (31 years with 1764 individual dive surveys). Surveys were conducted every year except for 2003 (due to budget cuts). Six sites were inside a no-take marine reserve, whereas the rest were in fished areas. The case-study was a useful test of our approach because the ecosystem is undergoing rapid change that was unprecedented in historical data, including climate warming, oceanographic change and species invasions (Ling and Keane 2024; Bates et al. 2014).

We included four species in our analysis (Table 1). These species were chosen out of >40 species in the dataset that were suitable for time-series analysis. They were chosen to provide contrasting abundance trends that could be used to test the new split test on extremes of abundance trends. The species exemplify increasing, collapsing, stable and fluctuating abundance trends (Fig. S1).

**Table 1.** The four species included in the analysis.

| Species names<br>(name for text) | Common name | Abundance<br>dynamics | Details |
| --- | --- | --- | --- |
| *Centrostephanus*<br>*rodgersii*<br>("increasing") | Long-spined sea<br>urchin | Increase | *C. rogersii* is a benthic grazer that can cause urchin barrens and has increased in abundance due to warming waters (Ling and Keane 2024) |
| *Tosia australis*<br>("collapse") | Southern biscuit star | Sudden unexpected<br>collapse | *T. australis* was a common reef seastar whose population unexpectedly collapsed, probably due to climate warming (Edgar et al. 2023) |
| *Jasus edwardsii*<br>("stable") | Southern rock<br>lobster | Stable abundance | *J. edwardsii* is an important fishery species and is a benthic predator that has stable abundance (Bates et al. 2014) |
| *Trachinops*<br>*caudimaculatus*<br>("fluctuating") | Hula fish | Stable but<br>fluctuating<br>abundance | *T. caudimaculatus* is a small fish whose abundance varies year to year likely due to changing ocean currents (Bates et al. 2014) |

For all training splits we used a timespan of 14 years, to ensure sufficient estimation of time-series parameters. We further divided survey sites into four regions for analysis (Fig. S2). These regions represent areas that share similar fishing pressure and recruitment patterns, so we treated sites within these regions as independent populations in the timeseries models.

The models included a principal component that represented changes in oceanographic conditions. This was built from a time-series of temperature, salinity, nitrate and silicate measured at the Maria Island National Marine Reference Station (Fig. S3, Lynch et al. 2014; CSIRO and IMOS 2024). The inclusion of covariates in the RW1 and AR1 models meant that intrinsic growth rates could be environmentally driven. The principal component was included in the model with 0, 1 and 2 year lags, and we used shrinkage priors to guard against overfitting.

We fitted and tested models for four species, 15 forecast horizons, 15 origins, two splitting rules and three different model structures. This resulted in 2800 combinations of treatments and 140,544 evaluations of forecasts at the survey level. Forecast evaluations have high variability (Ward et al. 2014; Coron et al. 2012) so we summarized trends in the test statistic by grouping results into three epochs: near (2007-2011), mid (2012-2017) and long-term (2018-2021) (fig. 2), and plotted the test statistic against the forecast horizon by each species and model. The error statistics were summarized by fitting then plotting GAM fits for each panel (Wood 2017).

Given our hypotheses we inspected the GAM fits to explore for trends of (1) forecast errors increasing for longer forecast horizons; (2) errors for the legacy splitting rule for epochs that were further from the training period would be larger than errors evaluated with the modern splitting rule; (3) the comparison of the legacy splitting rule to the modern splitting rule would show the biggest difference for species that had rapid changes in abundance, whereas the two splitting rules would give similar results for species that have little or gradual long-term change. Specifically, we expected that species that have recently expanded their ranges and increased (C. rodgersii) or decreased (T. australis) in abundance due to climate warming would have the biggest difference in the error statistic between the two splitting rules.

## Results

### Simulation testing

All models and both split-test rules had the same forecast errors for forecasts made from before the population collapse (fig. 3). Under simulations with a reduction in the carrying capacity there was a distinct forecast horizon at 10-15 years ahead, which coincided with the population collapse. This forecast horizon levelled out at a higher MASE for larger collapse levels (e.g. MASE of ~5 for a collapse to 25% of initial carrying capacity vs a MASE of ~2 for a collapse to 75% of initial carrying capacity).

**Figure 3.**
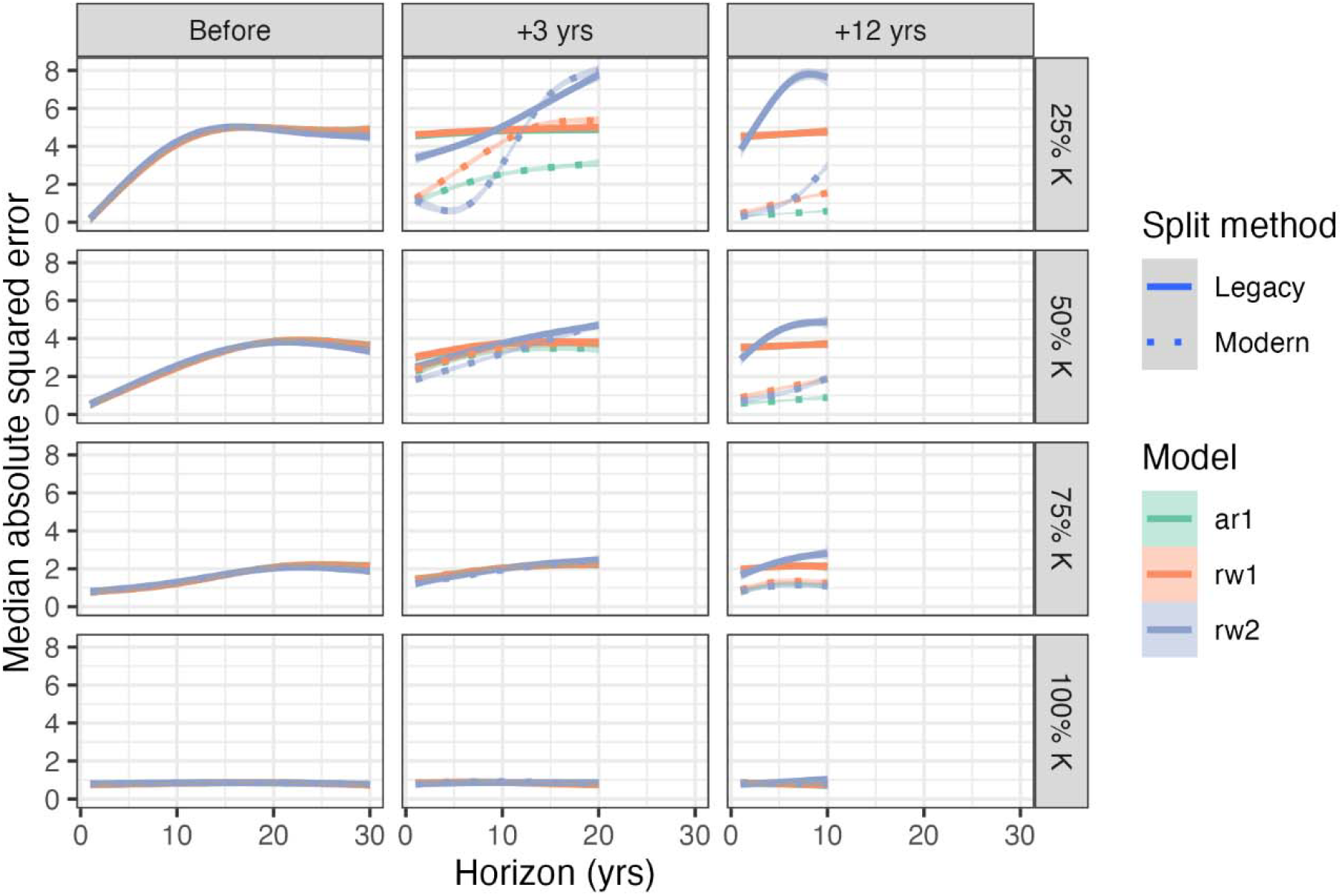
Results from simulation study showing forecast error for different time-horizons (number of years ahead of the origin) under the two split-test rules for a range of productivity collapses (% of carrying capacity, K) and for different forecast origins either before the collapse, 3 years after the collapse or 12 years after the collapse.

The legacy splitting rule performed as expected and estimated larger forecast errors than the modern rule when there were productivity declines in the forecast period. The largest differences in MASE for the split-test rules occurred when the forecast origin was well after the collapse and when the level of collapse was larger. For example, forecasts made forwards from 3 years after the collapse had a MASE of ~1.5 for all models with the rolling rule versus a MASE of 4-5 for the fixed training period rule (fig. 3). The RW2 model tended to have the worst performance under the legacy rule and large productivity declines, whereas it has similar performance to the RW1 under the modern rule.

### Case-study analysis

First we focus on results under the AR1 model (example forecasts in figs S4 & S5). The collapsing, stable and fluctuating species (*T. australis*, J. edwardsii and *T. caudimaculatus* respectively) had increases in error for longer time-horizons, consistent with our first hypothesis. The increasing species (*C. rodgersii*) had high errors at all epochs and that decreased slightly for longer time-horizons, a result that was counter to our first hypothesis (fig. 4).

**Figure 4.**
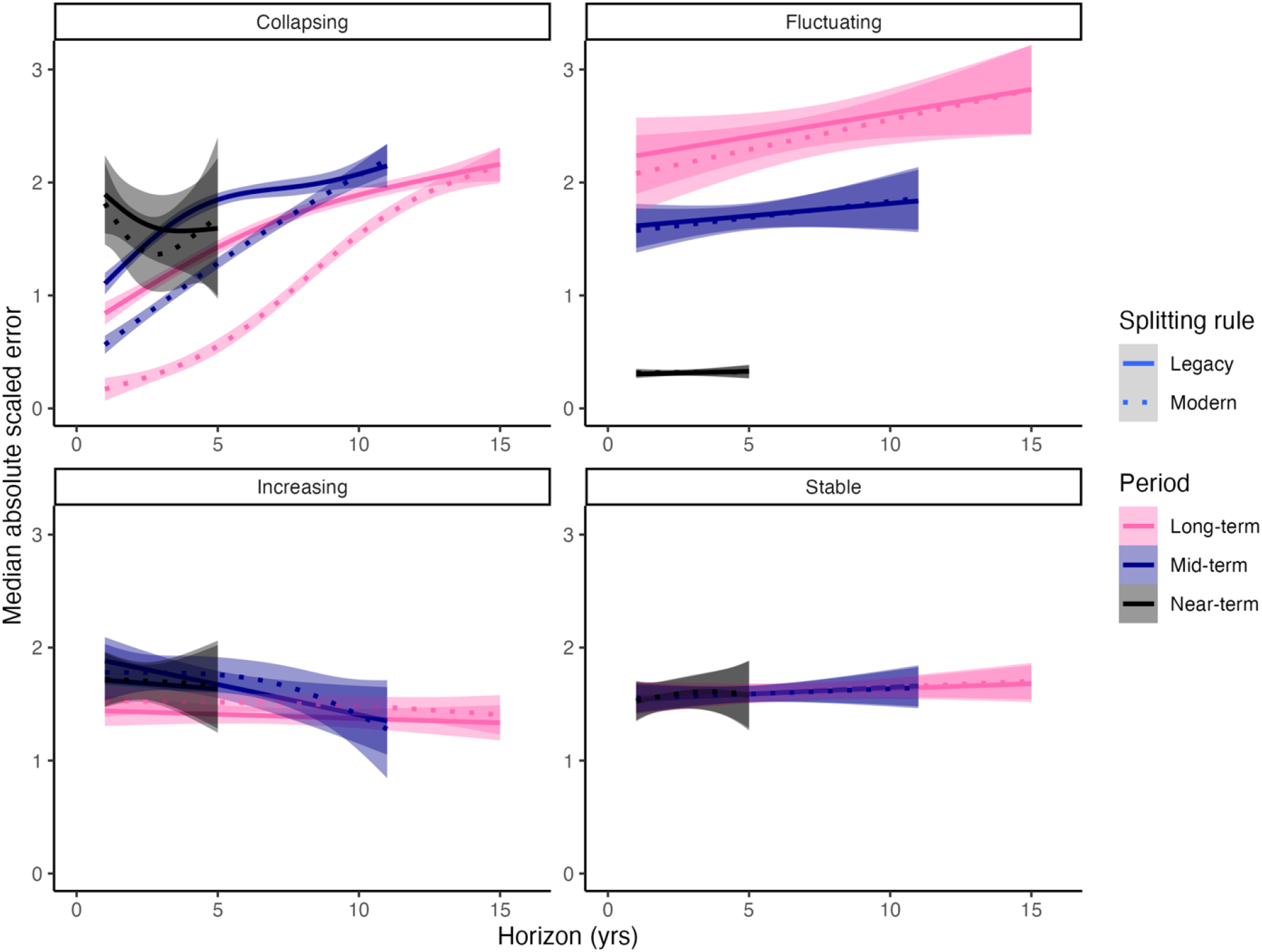
Comparison of forecast errors for the four case-study species, 15 time horizons and the two splitting rules. Results are shown for the AR1 model.

Results for hypothesis 2, that the legacy rule would have higher errors at more recent epochs when compared to the modern rule, were mixed. Errors for the stable species were the same regardless of the epoch, the collapsing species errors were greatest for the near-term epoch and lowest for the long-term epoch. For the fluctuating species errors were greatest for the long-term epoch and small for the near-term epoch. These patterns in forecast errors were not necessarily reflected in non-stationarity of parameter estimates under the rolling training period test rule with only the collapsing and increasing species showing temporal trends in parameter estimates (fig. 4).

The different patterns in forecast errors by species can be explained by their different abundance trends (figs 4 & S1). The collapsing species had a population collapse soon after the end of the initial training period (fig. S1), resulting in poor near-term predictions because forecasts were initialized in a period with high abundance. In later epochs abundances were near zero, so forecasts were initialized near zero and short time-horizon forecasts were more accurate than short time-horizon forecasts in the near-term epoch. For the long-term epoch, longer time-horizon forecasts had lower accuracy (because the model converged back towards the historical mean). The increasing and stable species had slowly increasing and stable abundances, respectively, so errors were similar at different epochs. The fluctuating species abundances went through large cycles, making longer-term and later epochs harder to forecast.

Only the collapsing species had worse forecast errors for legacy rule when compared to modern rule (hypotheses 2 and 3). The discrepancy between the two split-test rules was largest for the long-term epoch, consistent with the population collapse and non-stationarity in population parameters (Figure S6). Notably, the shape of the forecast horizon curve was different between split test rules, with the legacy rule having the steepest slope at a one year horizon, whereas the modern rule had a uniform slope (mid-term) or the steepest slope at intermediate time-horizons (long-term).

Only *T. australis* showed notably different results for the RW1 model when compared to the AR1 model (fig. S7). The errors under the RW1 were similar for the two split test rules. The default prediction for the RW1 is continuation of last year’s abundance, so the RW1 performed better than the AR1 when it was initialized post population collapse. The RW2 model tended to perform poorly, especially for longer time horizons or the long-term epoch (fig. S5). This model forecasts a continuation of the trend at the origin, so could make some particularly wild forecasts with high errors.

## Discussion

We observed that models fitted to legacy data could have poorer than expected predictive accuracy when forecasting collapsing species. Predictive accuracy with legacy data compared to modern data was poor because a model fitted to legacy data has parameter estimates that are no longer relevant in the contemporary environment. Our conclusion was further supported by the species with large inter-annual fluctuations but stationary population dynamics (*T. caudimaculatus*), having similar forecast errors under both rules. The collapsing species in our case-study, *T. australis*, may be indicative of the challenge of forecasting abundances in rapidly changing environments. Its sudden population collapse was unexpected and there is insufficient information about its biology to have anticipated this collapse, or to attribute its cause in hindsight (Edgar et al. 2023). The rapid environmental change at our case-study site is unprecedented in historical data at this location, and included ocean warming, oceanographic changes and species invasion (Bates et al. 2014; Ling and Keane 2024). All of these changes are plausible causes of the population collapse that ultimately contributed to lower forecast accuracy for this species.

The species that managers most need accurate forecasts for are those that are most likely to exhibit dramatic change in response to environmental changes, but our results show these species may also be the hardest to predict. Our finding provides a cautionary tale for forecasting abundances in other ecosystems: most of the world’s ecosystems are also facing no-analogue change (e.g. Williams, Jackson, and Kutzbach 2007) and for most biodiversity we lack sufficient information for assessing population trends (particularly invertebrates, Eisenhauer, Bonn, and Guerra 2019). These trends combined with our findings suggest that measurements of forecast accuracy can be too optimistic for future environments and have implications for the interpretation of ecological predictions by managers. For example, even if management was set to be appropriately conservative for known forecast errors, sudden population shifts can push populations well outside expected bounds of change.

Predictions made in time periods close to a collapse point (i.e. the near-term forecasts) or for species with stable or slowly changing population dynamics had similar accuracy for the legacy and modern split methods. There are two lessons in this result. First, not all species will exhibit dramatic population change and non-stationarity in parameters in response to rapidly changing environments. What is needed now is to identify how forecast errors vary across species and ecosystems, such as using species traits to identify generalities in forecast errors. Characterizing forecast errors can reveal insights that can improve ecological theory and management applications of forecasting (Ward et al. 2014; Lewis et al. 2023). For example, longer lived species tend to have lower forecast errors (Ward et al. 2014), a finding that could be used to set appropriate uncertainty boundaries for management, such as harvest limits (Raftery 2016), based on species life-history. Another past finding is that causal models can make more accurate forecasts than phenomenological models (Pichler and Hartig 2023)– this finding could be tested with our approach to see if the same is true under non-stationary dynamics. Split-test approaches that force no-analogue tests have been used for hydrological models to show good predictability for short duration droughts, but not for droughts of more than 3 years (Coron et al. 2012). The ecological equivalent would be to identify regions or species where our models are most likely to fail, which could then translate into more conservative management advice, such as on fishery harvest limits.

A second lesson was that forecasts based on legacy data but made for periods near to a collapse point will have similar accuracy to forecasts made with modern data. This result is complementary to the well-known increase in forecast errors for longer horizons (Petchey et al. 2015). The decay of forecast accuracy at longer time horizons is driven by multiple sources of uncertainty, including uncertainty in the initial condition, differences between system dynamics and modelled dynamics and environmental forcing. All of these factors contribute to the forecast horizon, a time forwards in the future at which predictions are too inaccurate to be useful for a stated purpose (i.e. errors cross a predefined maximum threshold) (Petchey et al. 2015). Comparisons of the legacy and modern split-test methods can inform on how quickly a fitted model passes its use-by-date. Counter-intuitively, we found for a collapsing species that predictions for the near-term epoch had higher errors than predictions for the mid and long-term epochs. This counter-intuitive result highlights the relative importance uncertainty in initial condition and uncertainty in system dynamics for forecast accuracy (Dietze 2017). Forecasting a few years ahead accurately near to a rapid state (abundance) change is harder than forecasting initialized well after the state change, even if the model parameters are less relevant at the longer time period. Forecast errors further from the forecast origin become more dominated by system uncertainty than by initial condition uncertainty (Dietze 2017), so legacy models will have relatively poorer performance at longer time horizons than modern models.

Forecast errors usually get broader the further ahead of the forecast origin we are predicting. Unusually, errors for our increasing species (*C. rodgersii*) decreased slightly for times further in the future. This species is undergoing a rapid population expansion in Tasmania, as it benefits from warming waters (Ling and Keane 2024). Over time, its abundance dynamics are stabilizing, though high density urchin barren formation still remains patchy in the area of our study. Thus, its invasion and population expansion is becoming increasingly certain.

The split-test approach we used is based on observational data, so is potentially subject to confounding, such as between time period and the number of years forward from the origin. Our approach is also retrospective, and only those species that historically had large shifts in abundance will be characterized as having high forecast errors. Our approach does not provide any evidence for or against other species having similar sudden shifts in the future. An alternative would be to use simulated data from a ‘known’ model that represents the truth. We chose not to rely fully on simulated data here, because it does not represent the true complexity of timeseries data (Storch et al. 2017). Further, split-test validation is a common approach across many disciplines of timeseries analysis (Tashman 2000). In other disciplines, split-test validation has already proven to provide advances in forecasting skill, by helping reveal generalities in the types of environments that will be hardest to forecast (Coron et al. 2012). Therefore, we recommend more ecological studies use split test validation approaches to advance our knowledge of when and for what types of species forecasts will be most reliable.

In some cases, forecast errors could be improved by using more complex models. For example, our models did not account for multiple species interactions. The case-study system is controlled by strong ecological interactions, such as the role of lobster as ecosystem engineers that prevent urchin barren formation (Bates et al. 2014; Ling and Keane 2024). It is likely that a model that accounted for species interactions could improve forecast skill. Future studies could explore approaches such as state space structural equation models (Brown et al. 2023) or hierarchical multispecies models (Ovaskainen et al. 2017). Comparison of forecast skill for multispecies and single species would then provide evidence for or against the role of species interactions in driving abundance dynamics. We additionally predict that multispecies models will perform better when extrapolated to no-analogue futures, because they better account for ecological dynamics. The goal of reducing errors through enhanced modelling should be pursued concurrently to the goal we addressed here of characterizing model errors, because, ultimately, the ability to reduce forecast errors is limited by the complexity of ecosystems and difficultly of making accurate measurements (Beckage, Gross, and Kauffman 2011; Dietze 2017).

## Conclusion

We suggested a new approach for splitting time-series to quantify forecast errors. Our approach reveals that forecast errors can dramatically worsen in situations where there are rapid changes in population abundance. Appropriate quantification of forecast errors is important to ensure management decisions account for uncertainty, and as we seek to use forecasting to test competing ecological theories. As humanity continues to change the earth system outside the bounds of historical observations, our ability to predict change will likely worsen. Our new approach provides a way to quantify and communicate how much worse our predictions may be.

## Author contributions

CJB, CAB and RSS conceived the ideas and designed methodology; NB, RSS, GJE and LO collected the data; CJB analysed the data; CJB led the writing of the manuscript; all authors contributing interpreting the results, designing visualizations, and writing and editing the manuscript. Statement on inclusion: our study includes scientists working with ecosystem managers in the case-study region.

## Acknowledgements

CJB was supported by a Future Fellowship (FT210100792) from the Australian Research Council. The National Marine Reference Station dataset was collected by CSIRO and multiple contributors including field and laboratory staff involved in the CSIRO Coastal Monitoring program from 1944 - 2008. Val Latham CMAR 1991-2000’s (Hydrological analyses) Integrated Marine Observing System (IMOS) April 2009 –

## Data availability

Biological data from the Australian Temperate Reef Collaboration and Reef Life Survey are available at: https://portal.aodn.org.au/ (Edgar and Stuart-Smith 2014; Edgar and Barrett 2012). The ecological data used for this study are managed through, and were sourced from, Australia’s Integrated Marine Observing System (IMOS) – IMOS is enabled by the National Collaborative Research Infrastructure Strategy (NCRIS)

Physical oceanographic data are available at: https://www.marine.csiro.au/data/trawler/ (Lynch et al. 2014)

