## Supplemental methods and figures for "Assessing predictive accuracy of species abundance models in dynamic systems"

### Supplementary material

**Supplemental methods**

*Implementation of the legacy splitting method*

The legacy splitting method required recalibrating model parameters. To recalibrate parameters, we first trained the model on the training period. We then fixed the hyperparameters for random effects and fixed effects at the values from the initial fit and refit the model’s random effect estimates with data up until each forecast origin.

We chose to use Bayesian time-series models because they allowed us to model timeseries of counts with an appropriate likelihood (poisson or negative binomial for counts). The R-INLA program is also very computationally efficient, so facilitates the repeat fitting required by cross-validation. INLA provides flexible model structures that include autoregressive and random walk models (Rue et al. 2017). Further, timeseries models of populations can be hierarchically structured to allow for multiple regions (Brown and Roff 2019). A final advantage of the Bayesian model is that it is straightforward to make forecasts and fill missing values.

The implementation of INLA meant it was practical to implement the recalibration of model parameters for the legacy split method. The recalibration was necessary so that we could forecast from a new origin, but using parameters that were from data well before the origin. This is best explained with an example. Say we wanted to apply the legacy method with training data from 1992 to 2005, but initialized the forecast origin at 2015. We would first fit the model to data from 1992-2005. Then we would create a new INLA model where the hyper-parameters (e.g. rho) and fixed effects (intercept, environmental effects) were given priors centered on the values estimated from the first fit. The precisions for these parameters were set 10^12^, effectively forcing constant parameter values. The model was then refitted on data from 1992 to 2015. Thus, the random effects (ie year-by-year process noise variations in the AR1) were calibrated up until 2015, which initialized forecasts forward of 2015 at the 2015 value. INLA makes predictions (forecasts) at the same time as it fits, so data for 2016 to 2022 were included as NA values to allow for predicting without fitting. We could then compare the forecasts from 2016 to 2022 to the observed values, which were held back.

*Further details of the principal components analysis of environmental variables for the case-study*

The environmental covariates were measured approximately monthly, but not consistently, so we needed to standardize for unequal sampling intervals before their use in the principal components analysis. For each environmental variable, we fitted generalized additive models (GAMs) with interannual and seasonal splines (Wood 2017). Means from the model’s predictions for January where then used in the principal components analysis. The principal components were designed to be representative of changes in currents and environmental conditions that may impact productivity and recruitment of our species (PC1 – 46% variation, Fig. S3).

Wood, Simon N. 2017. *Generalized Additive Models: An Introduction with R*. chapman and hall/CRC.

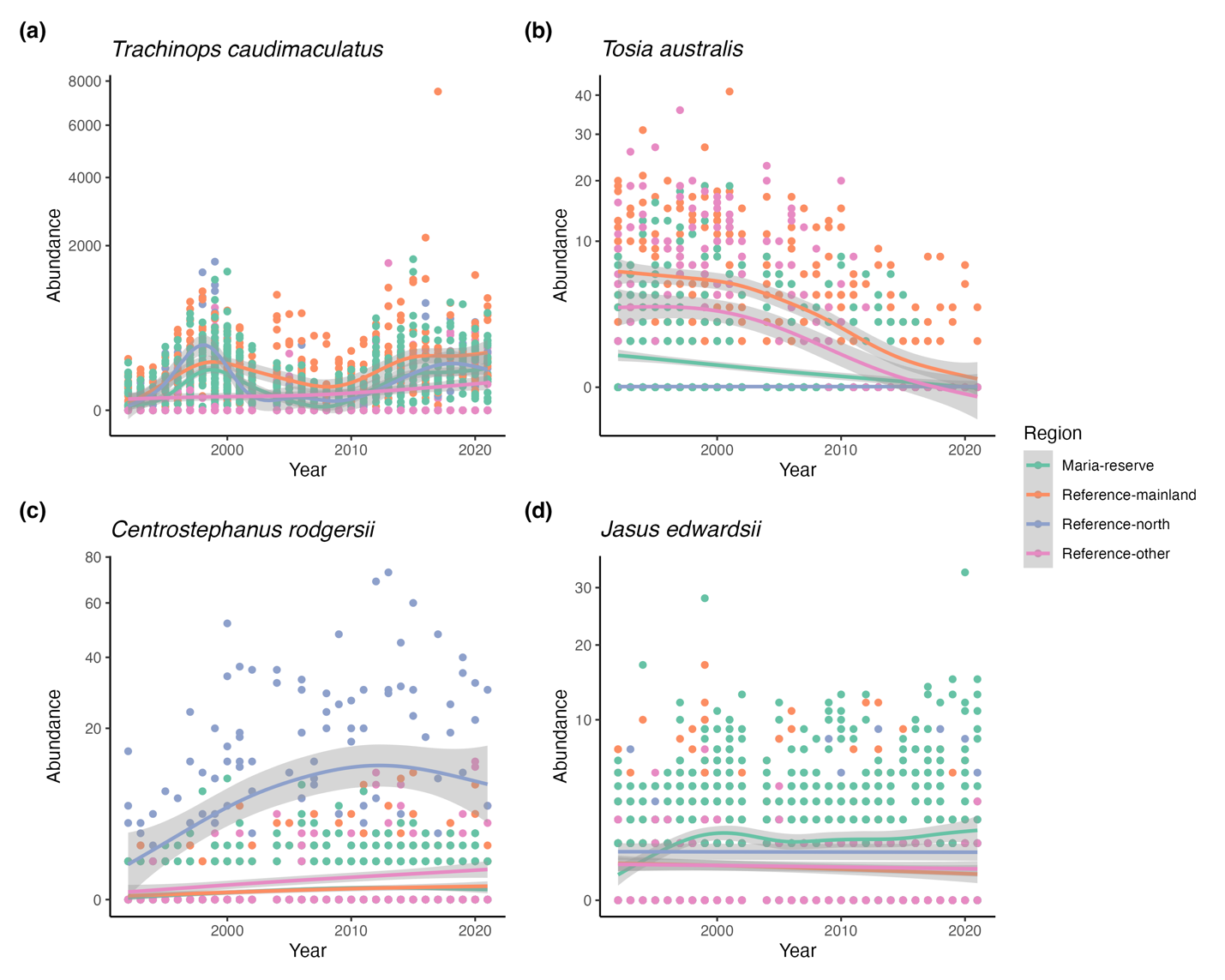

**Figure S1** Abundance data with spline trends by regions for each of the case-study species.

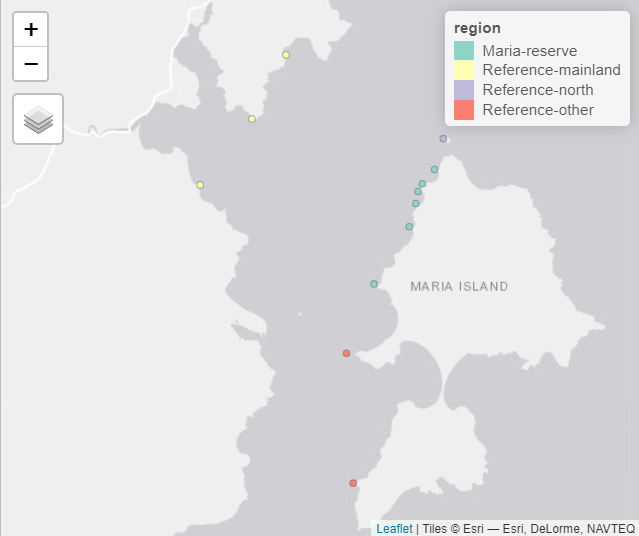

**Figure S2** Map of study sites and their division into regions for modelling

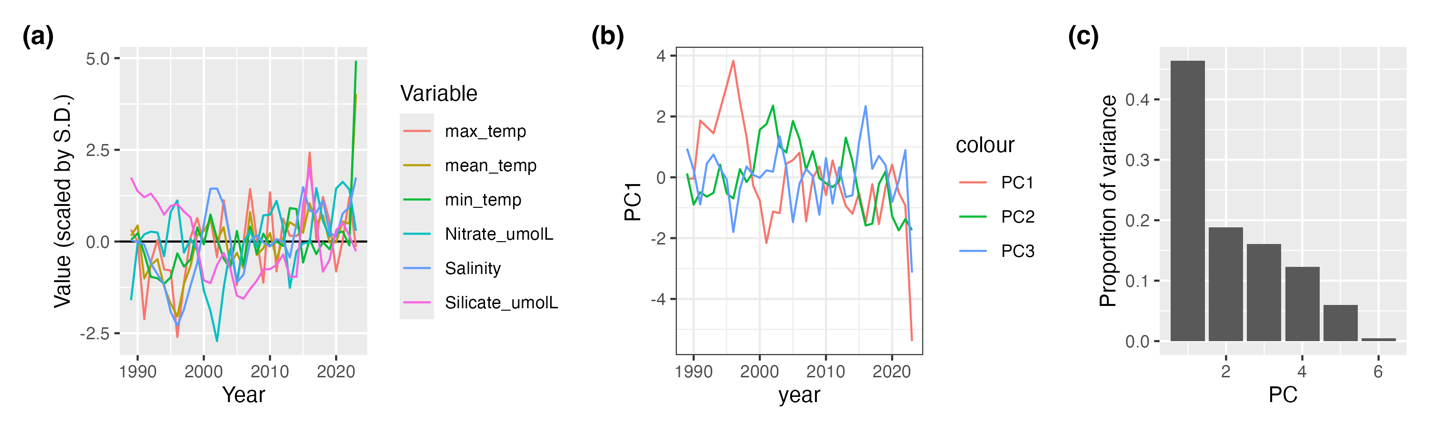

**Figure S3** Covariate timeseries, principal component time-series and variance explained by each principal component.

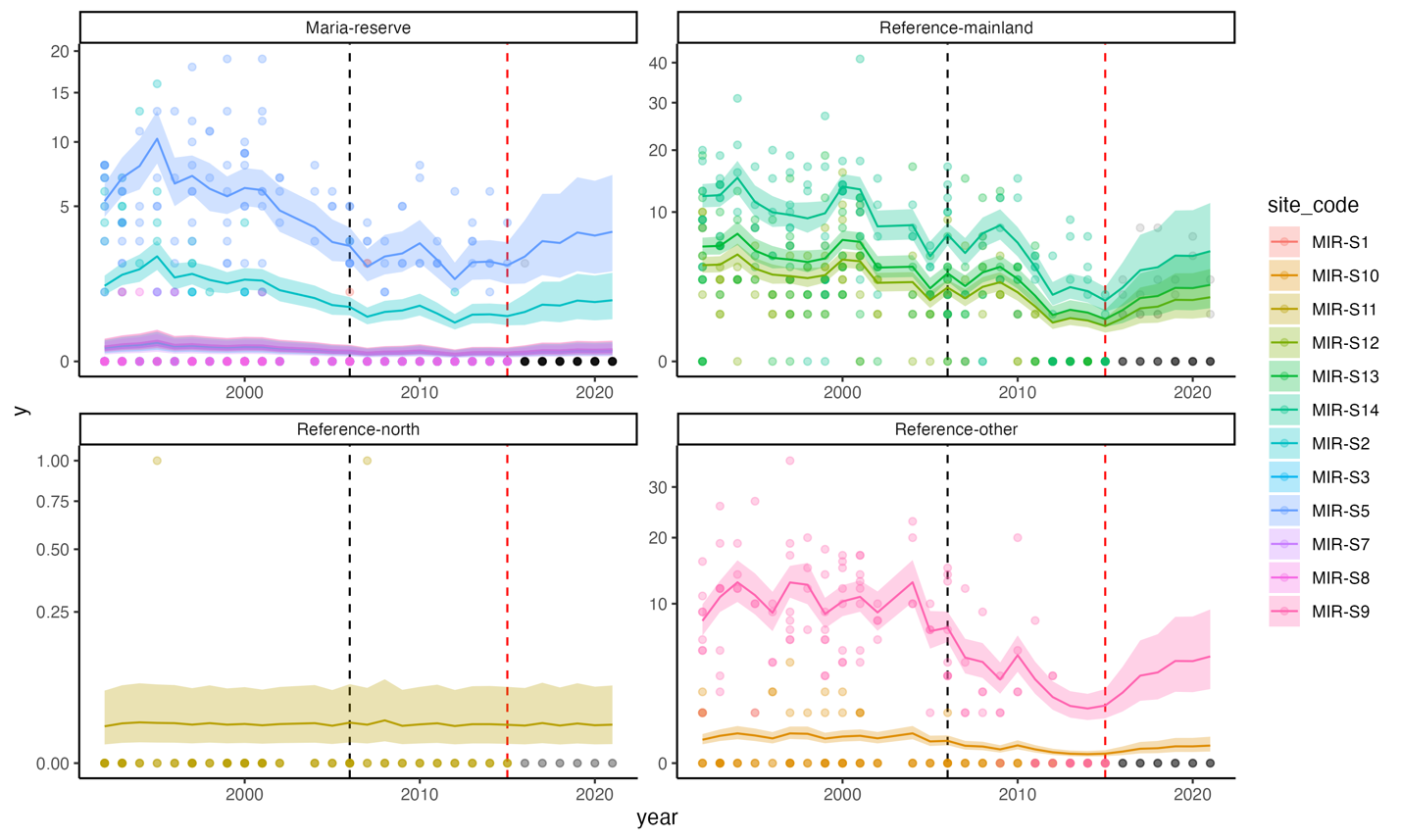

**Figure S4** Example fits and forecasts for *Tosia australis* from the ar1 model with the legacy split method and a forecast origin of 2015. Black dashed line shows the final year of training, the red dashed line shows the forecast origin. Lines show median predictions for each site and shaded regions show 95% credible intervals. Points show observed abundances.

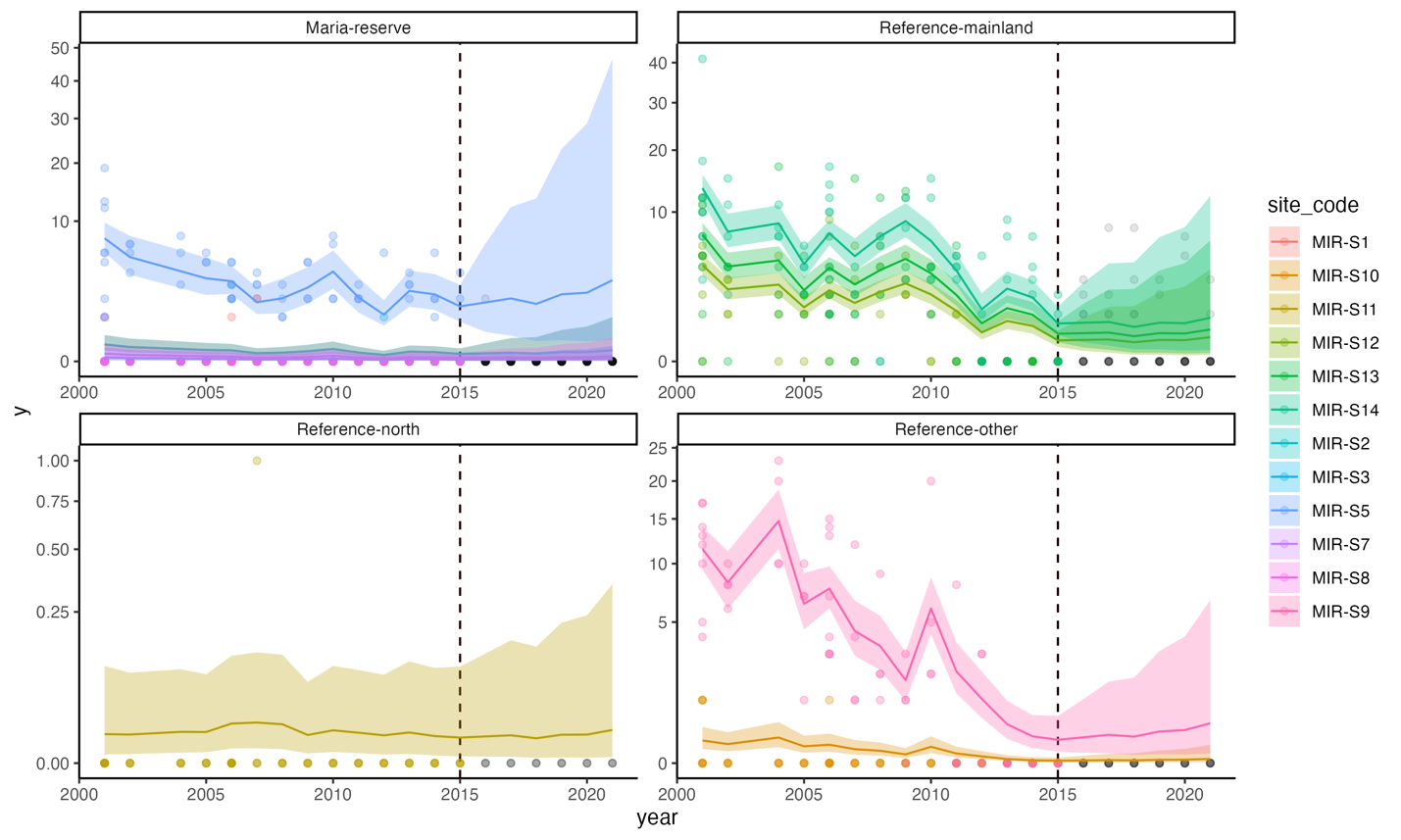

**Figure S5** Example fits and forecasts for *Tosia australis* from the ar1 model with the modern split method and a forecast origin of 2015. Black dashed line shows the final year of training which is also the forecast origin. Lines show median predictions for each site and shaded regions show 95% credible intervals. Points show observed abundances.

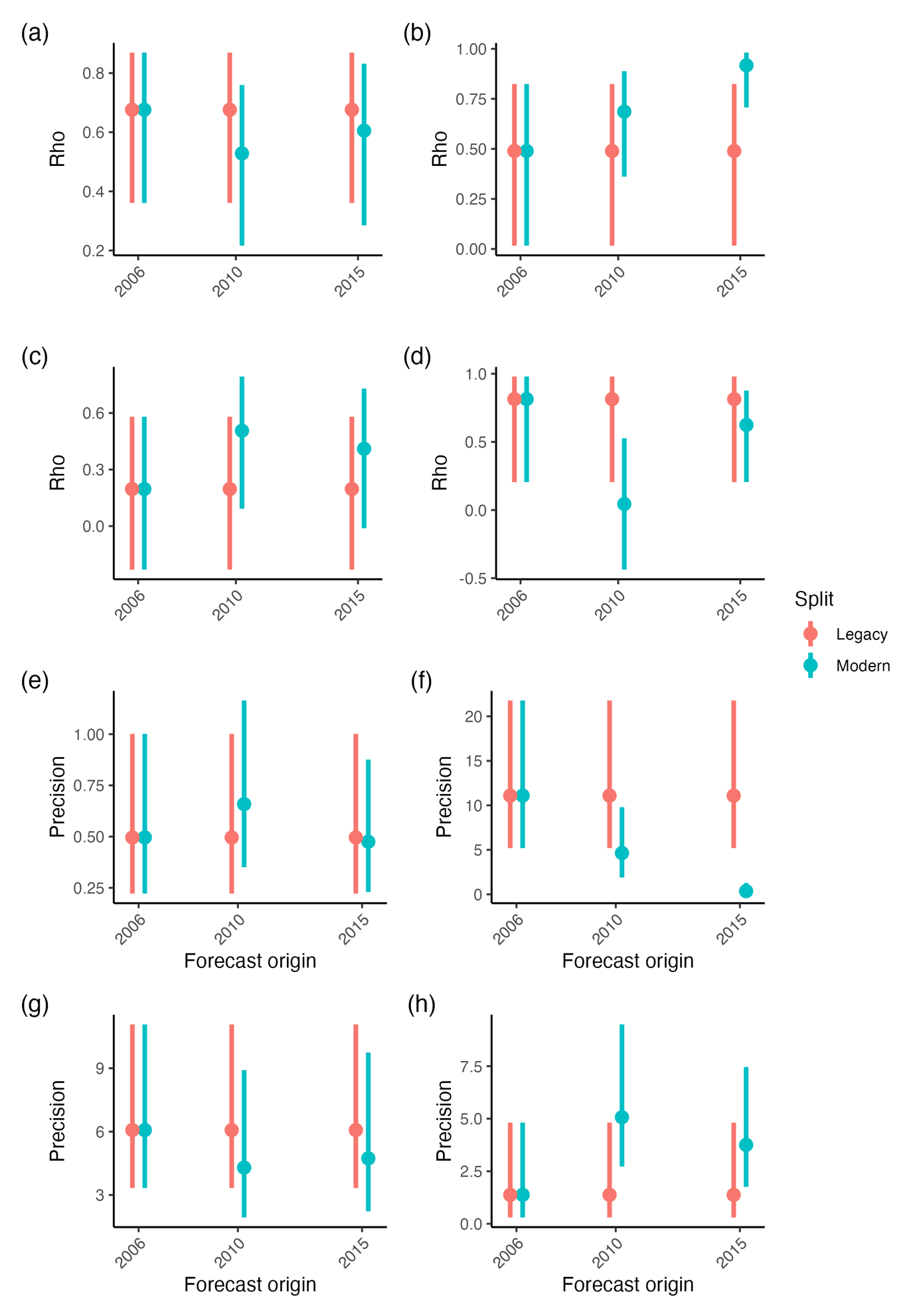

**Figure S6** Estimates for $\rho$ (a-d) and precision of inter-annual year variation (e-h) for each split-test approach and forecast origin. Panels show *Trachinops caudimaculatus* (a and e), *Tosia australis* (b and f), *Jasus edwardsii* (c and g) and *Centrostephanus rodgersii* (d and h). Points show median, bars show 95% credible intervals.

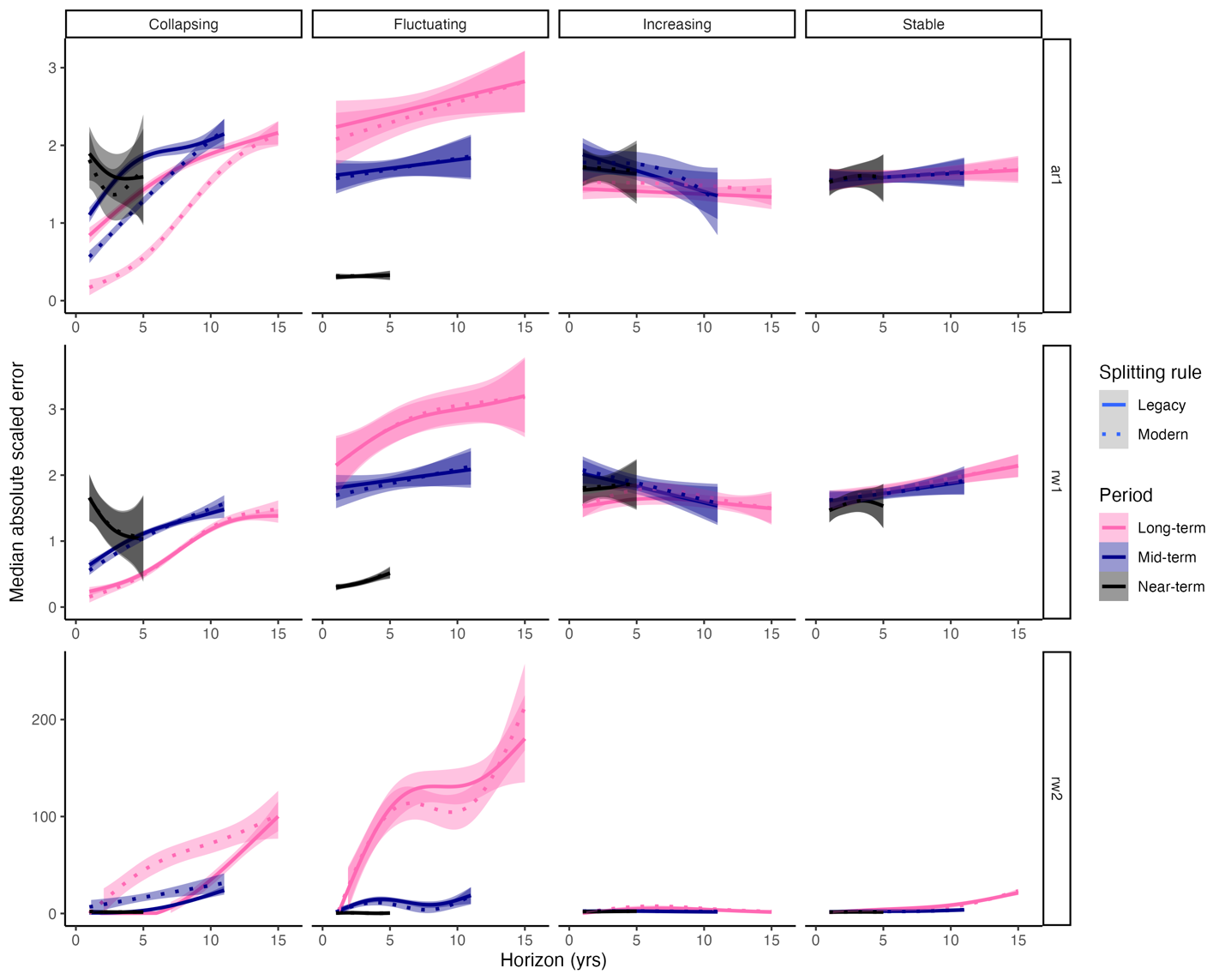

**Figure S7** Forecast errors for both splitting rules, all species and all models. Note that MASE values were truncated at 1000 for plotting.

**Table S1** Models fitted and their interpretation

| Model name (‘short name’) | INLA Formula | Population dynamics assumption | Explanation |
| --- | --- | --- | --- |
| Random walk order 1 - reference model (RW1) | y ~ 1 + f(site_code, model = “iid”) + f(year, model = “rw1”, replicate = iregion)+ f(enviro, model = “linear”, mean.linear = 0, prec.linear = 1) | No density dependence, one population per region, populations share intrinsic growth parameters. Allows for environmental effects on the intrinsic growth rate (Brown and Roff 2019) | Abundance within each region follows an independent random walk (regions shared the prior and variance parameter ). The multiplicative effect of environmental covariates on growth rate is shared across populations. |
| Autoregressive order 1 model(‘AR1’) | y ~ 1 + f(year, model = “ar1”, replicate = iregion) + f(region, model = “iid”) + f(site_code, model = “iid”) + f(enviro, model = “linear”, mean.linear = 0, prec.linear = 1) | Density dependence, one population per region, populations share intrinsic growth parameters but have independent carrying capacities. Allows for environmental effects on the intrinsic growth rate (Ross et al. 2015) | Abundance within each region follows an independent autoregressive process. (regions shared the priors and variance and autocorrelation parameters - meaning we assumed the different regions had the same growth and carrying capacity parameters). The multiplicative effect of environmental covariates on growth rate is shared across populations. |
| Second order random walk (‘RW2’) | rw2_site = mref_form <- y ~ 1 + f(site_code, model = “iid”) + f(year, model = “rw2”, replicate = iregion, constr = FALSE, scale.model = TRUE, hyper = prec.prior)+ f(enviro, model = “linear”, mean.linear = 0, prec.linear = 1) | Not a population dynamics model, assumes continuation of trend (smoothed). Allows for multiplicative environmental effects on mean abundance | Abundance at each site follows an independent second order random walk (shared variance parameter). Environmental effects shared across regions. |

**Table S2** Parameters used in the simulation study

|  | **Parameter** | **Value** | **Explanation** |
| --- | --- | --- | --- |
|  | Time-series length | 55 | Long enough to give stable results in simulation tests with only single time-series, but short enough to be realistic for ecology |
|  | Year of productivity change | 32 | Chosen to enable sufficient data for testing pre and post regime shift forecasts |
|  | Training series length | 25 | Long enough to enable stable time-series model fits for simulation testing |
|  | Productivity changes | 0.25x, 0.5x, 0.75x and 1x initial carrying capacity at year 32 | Chosen to represent a range of magnitudes |
|  | Number of time-series simulated per productivity change value | 20 | Chosen so as large enough to get stable results, but small enough for computational efficiency (for each unique time-series we then need to fit and test six models: two split methods by three training and testing epochs by four productivity changes, so 20 replicates requires 480 model refits) |
| Epochs for plotting | | Years 25 (7 years before productivity change), 35 (3 years after productivity change) and 45 (12 years after productivity change) | Chosen so as to provide results before and after the productivity change |
|  | Time-series model for fitting | AR1 | Used the AR1 as it represents a best-case scenario where the fitted model is the same as the model used for simulation. |
|  | Gompertz model initial abundance | 20 | Set at carrying capacity |
|  | Gompertz growth rate | 0.25 | Realistic mid-range value |
|  | Gompertz carrying capacity | 20 | Arbitrary |
|  | Process noise S.D | 0.05 | Chosen to give realistic levels of population size variation |
